# *Atxn2*-CAG100-knock-in affects mouse lifespan and vestibulo-cerebellar function via neural disconnection

**DOI:** 10.1101/333443

**Authors:** Melanie V. Halbach, Nesli-Ece Sen, Júlia Canet-Pons, Bram W. Kuppens, Mandy Segers, Martijn Schonewille, Ewa Rollmann, Kay Seidel, Udo Rüb, David Meierhofer, Michel Mittelbronn, Patrick Harter, Chris I. De Zeeuw, Luis E. Almaguer-Mederos, Suzana Gispert, Laurens W.J. Bosman, Georg Auburger

## Abstract

Unstable expansions in the Q22-polyglutamine domain of human ATXN2 mediate risks for motor neuron diseases such as ALS/FTLD or cause the autosomal dominant Spinocerebellar Ataxia type 2 (SCA2), but the pathogenesis is not understood and models are unavailable.

We generated a novel knock-in mouse line with CAG100 expansion in *Atxn2*, transmitted unstably. The mutant protein accumulated in neuronal cytosolic aggregates, with a characteristic pattern of multi-system-atrophy. Loss-of-function phenotypes included less mutant offspring, initial weight gain and motor hyperactivity. Progressive toxic aggregation effects started around 20 weeks in homozygous animals showing weight loss, reduced muscle strength and gait ataxia. Lifespan was decreased. In the cerebellum, neuronal soma and dendrites were remarkably spared. However, myelin proteins MBP, CNP, PLP1 and transcripts *Mal, Mobp, Rtn4* decreased markedly, especially adhesion factors MAG and MOG. In neurons, strong reductions were found for mRNAs of perineuronal elements *Haplnl, Hapln2, Hapln4*, of axonal myelin interactors *Prnp* and *Klk6*. At protein level, the adhesion factor neuroplastin and neurofilaments were strongly reduced, while presynaptic alpha-synuclein increased two-fold.

Overall, this authentic SCA2 mouse model elucidates how altered function and aggregation toxicity of ATXN2 conspire to trigger axon-myelin disconnection. This model will promote the development of neuroprotective therapies and disease biomarkers.

## Introduction

During periods of stress, for example as a consequence of starvation, RNA processing, quality control and trafficking, as well as translation, have to be adapted to adjust cell metabolism to the adverse conditions. This is especially important for the survival of neurons as they are particularly sensitive to stress. Tightly controlled by regulatory proteins such as Ataxin-2 (ATXN2), neurons depend on the selective production of a myriad of protein isoforms for the extracellular space, their membranes and their mitochondria to cope with stressful situations. ATXN2 and Ataxin-2-like (AXTN2L) represent a gene family with strong conservation, of which every eukaryotic species has at least one member, such as PBP1 in yeast. Most vertebrates and land plants have at least two orthologs, including *ATXN2* and *ATXN2L* in man and mouse, and CID3 and CID4 in *A. thaliana* (1). Family members have in common firstly a PAM2 domain that allows them to interact with the poly(A) binding protein (PABP), thus influencing the 3’-untranslated region of mRNAs during translation. Secondly, they share LSM domains that allow them to perform RNA processing tasks. Thirdly, they exhibit proline-rich motifs that modulate trophic receptor endocytosis and growth pathways (2–4).

Under normal conditions, ATXN2 is located in the cytosol, mostly associated with the ribosomal machinery at the endoplasmic reticulum (ER), where it modulates protein synthesis at the mRNA translation step (5, 6). Upon cell damage or bioenergetic deficits, its transcription is enhanced and ATXN2 is re-localized to stress granules (7), where mRNAs are stalled to undergo quality control until protein synthesis is resumed (8). Strong selective effects of ATXN2 orthologs on mitochondrial precursor transcripts were observed, in particular the leucine homeostasis factor IVD and the stress-response kinase PINK1, where mutations can cause Parkinson’s disease (9–12). In interaction with specific proteins like TDP-43 and ITPR1, ATXN2 acts as modulator of RNA splicing and neural excitability (13, 14).

The knock-out of ATXN2 in mouse triggers phenotypes of obesity, dyslipidemia, insulin resistance, with hepatic accumulation of lipid droplets and glycogen (4). In man, the chromosomal *ATXN2* locus was linked to obesity, hypertension and diabetes mellitus type 1 (15). Intriguingly, ATXN2 deficiency not only rescues the lethality of yeast PABP deletion (16), but also shows therapeutic relevance by mitigating the neurodegenerative process in spinocerebellar ataxias and motor neuron disease (17–19).

Conversely, phenotypes of neuronal atrophy were documented when ATXN2 contains unstable expansions of its polyglutamine (polyQ) domain that render it more insoluble and aggregation-prone. This domain has Q22 in man, but Q1 in mouse, encoded by CAG. ATXN2 was named by its initial description as the gene mutated in autosomal dominant Spinocerebellar Ataxia type 2 (SCA2), where patients carry pure (CAG)-repeat expansions to sizes ≥33 (20). Larger expansions lead to younger onset ages, faster progression, more widespread pathology and earlier death (21–23). Patients with repeat size Q92 and Q116 had clinical manifestation within the first year of life and showed multi-system atrophy of cerebellum, cerebrum, and brainstem (24, 25). When an intermediate expansion to 29-30 triplets occurs, usually with one remaining CAA interruption, it increases the risk to develop motor neuron diseases like ALS (Amyotrophic Lateral Sclerosis) or FTLD (Fronto-Temporal Lobar Dementia) (13). Also a specific haplotype of single nucleotide polymorphisms in *ATXN2* is associated with the risk of ALS (26). In addition, small expansions of ATXN2 with CAA interruptions were reported to underlie dopaminergic midbrain neuron atrophy in families with Parkinson’s disease (PD) (27).

Currently, there is no curative treatment for SCA2. Recent pharmacological and genetic developments open new opportunities in the near future. Although several models for SCA2 were developed, an authentic mouse model was still lacking. Previous mouse models largely focused on the overexpression of expanded ATXN2 in Purkinje neurons, so they are unsuitable to study extra-cerebellar deficits or the contribution of partial loss-of-function effects. Analysis of these mouse mutants showed that aggregates of ATXN2 protein accumulate in the cytosol rather than in the nucleus (28). Q58- and Q127-ATXN2 expansions alter neuronal excitability (14, 29–31). More recent efforts overexpressed a human Q72-ATXN2 BAC clone containing the physiological promoter and exon-intron structure and showed the G protein signaling factor RGS8 to be dysregulated in cerebellum (18, 32). We recently published the first knock-in (KIN) mouse where a CAG42-expansion triggers ATXN2 to sequestrate PABP into insolubility in vulnerable brain regions (33). An induction of the ubiquitination enzyme FBXW8 was observed as a cellular effort to degrade Q42-expanded ATXN2. In addition, partial loss-of-function effects dysregulate calcium homeostasis factors in these mice. Unfortunately, the phenotypes appear only after two years in these Atxn2-CAG42-KIN mice making them unsuitable for studying advanced stages of SCA2.

Here we present a new mouse model that shows authentic SCA2 with fatal course, created via knock-in of the clinically relevant repeat length of 100 CAG-trinucleotides into the murine *Atxn2* gene with intact murine promoter and exon-intron structure, in order to preserve its expression regulation.

## Results

### Generation of the *Atxn2*-CAG100-KIN and its stability

In order to study the progression of neurodegeneration at multiple levels in an authentic rodent model for SCA2, we created the novel *Atxn2*-CAG100-knock-in (KIN) mouse line. To this end, a (CAG)_100_ repeat with neighboring sequences was synthesized and inserted at position Q156 into the murine *Atxn2* exon 1 with flanking loxP sites, employing the homologous recombination strategy shown in Fig. S1A and using previously described targeting vectors (4, 33). Embryonal stem (ES) cell lines with successful knock-in underwent Flp-mediated excision of the neomycin resistance cassette and verification of the expansion length in heterozygous (CAG1/CAG100) or homozygous (CAG100/CAG100) state by Sanger sequencing and by PCR with Neo-flanking or repeat-flanking primers (Fig. S1B/C, Table S1), resulting in a single *Atxn2*-CAG100-KIN mouse line.

Genotyping with repeat-flanking primers and with DNA-fragment-length-analysis on polyacrylamide gels initially suggested stability of the expansion size, as previously observed in the Atxn2-CAG42-KIN mice (33). The expanded size of *Atxn2* mRNA was confirmed by saturation reverse-transcriptase (RT)-PCR in the cerebellar transcriptome from heterozygous (CAG100Het) and homozygous (CAG100Hom) knock-in mice (Fig. S1D). The expression level of the expanded *Atxn2* mRNA was slightly, but not significantly (p= 0.09) reduced in cerebellum at the age of 20 weeks (Fig. S1E). Thus, our new mouse mutant recapitulates the repeat expansion of patients at the DNA and the RNA level.

Next, we studied the abundance and solubility of the expanded ATXN2 protein. While WT and the Q42-expanded ATXN2 were soluble in the mild detergents containing RIPA buffer, two antibodies failed to detect the Q100-expanded ATXN2 among RIPA-soluble proteins of cerebellum at the age of 14 months (Fig. S1F-G). This included analyses with the monoclonal anti-polyQ antibody 1C2, which was raised against a 38Q motif in the N-terminus of nuclear TATA-box-binding protein and recognizes also other expanded polyQ domains, as well as analyses with the monoclonal anti-ATXN2 antibody. The Q100-ATXN2 was weakly detectable, however, in the more insoluble SDS-fraction of cerebellar tissue from CAG100Het and CAG100Hom mice (Fig. S1H). We take this as evidence that Q100-ATXN2 has a strongly reduced solubility in the cytosol compared to the wildtype protein.

Upon analysis of non-neural peripheral cells, the Q100-expanded ATXN2 was readily detectable by immunoblots of RIPA-soluble proteins from homozygous murine embryonal fibroblasts (MEF), although the amount of expanded ATXN2 corresponded to only 9% of wildtype (WT) (Fig. S1I). There was no Q100-ATXN2 in the SDS fraction. Immunocytochemistry of MEF showed a diffuse cytosolic distribution of ATXN2 without aggregation under normal conditions, and its presence in stress granules after arsenite exposure (Fig. S2). These observations indicate that the CAG100-expansion reduces the total protein levels of ATXN2. Q100-ATXN2 solubility appears to be lower in cerebellum than in MEF, compatible with the notion that the protein acquires an altered conformation in cells with strong excitability such as neurons. This seems credible in view of the excitation-induced aggregation of polyQ-expanded Ataxin-3 in neurons from patients with Machado-Joseph disease (34).

In patients, repeat expansions often increase over generations and show a mosaic pattern within somatic cells. To our knowledge, this instability and mosaicism has not been observed in previously generated mouse models of SCA2, while it is a known feature of mice with polyglutamine expansion that model Huntington’s disease (35). Although genotyping had initially suggested repeat stability in our Atxn2-CAG100-KIN mice, periodic testing in successive generations of this colony indicated the occurrence of further expansions and somatic mosaicism, as in SCA2 patients (Fig. 1).

**Figure 1:**
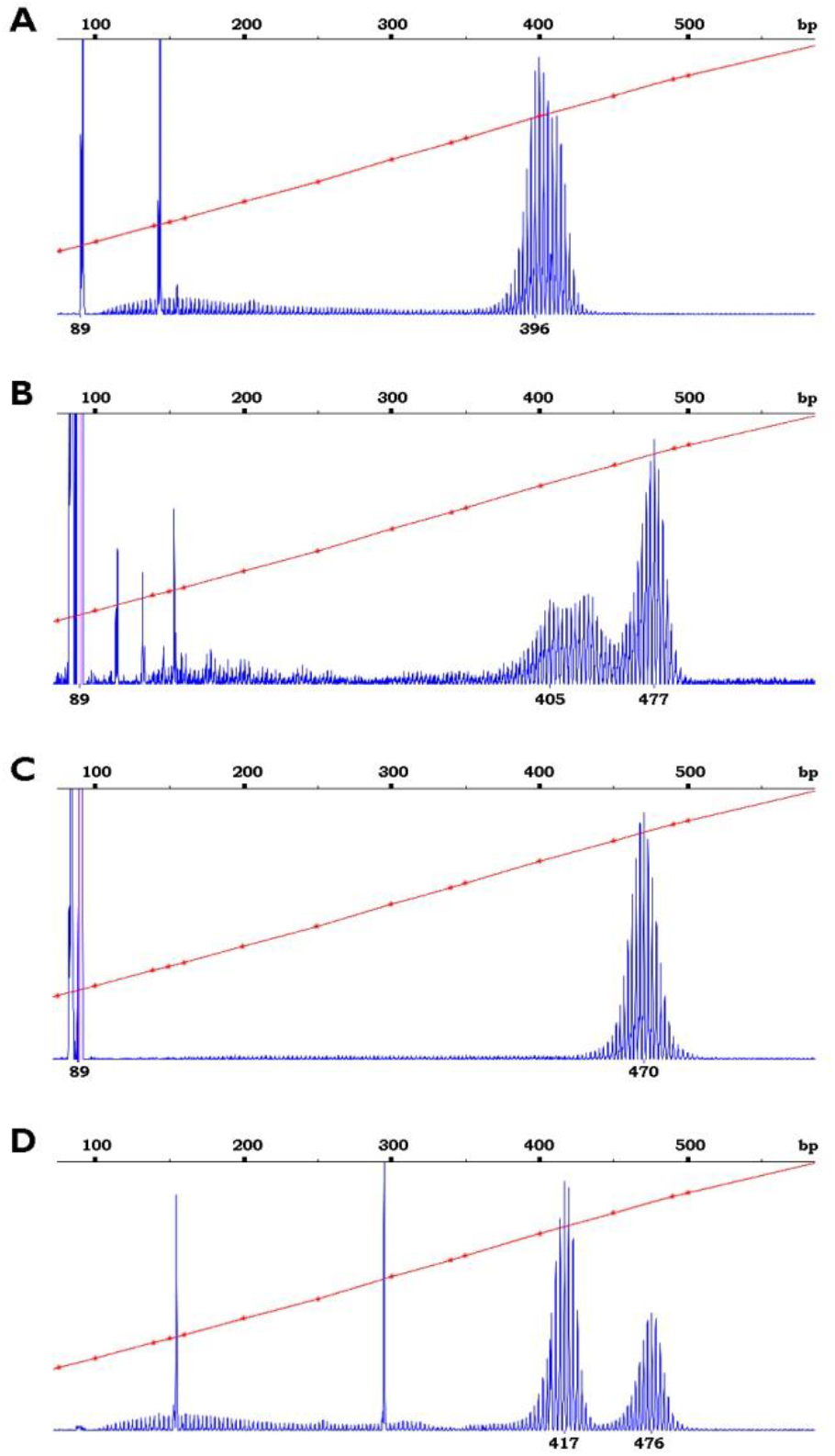
Instability and somatic mosaicism of CAG-repeat size, with expansion over successive mouse generations. Fragment length analysis of the CAG-repeat revealed instability of the CAG100 allele [predicted PCR fragment length of 387 basepairs (bp)]. DNA was retrieved from ear punches, amplified by PCR, loaded on polyacrylamide gels and compared to fluorescent size markers (diagonal lines). In CAG100Het mice, a sharp peak representing a PCR fragment length of 90 bp corresponded to the WT CAG1 allele **(A-C)**, while the mutant allele of larger size showed a hedgehog-like pattern reflecting somatic mosaicism **(A-D)**. Panels A-D show mice from different generations in chronological order, revealing an increase of the repeat length over a period of 4 years. To improve visibility, the peak representing the WT allele was clipped. In the CAG100Het mouse of the first lane **(A)**, the expanded allele showed an average size of 396.81 (the mosaic ranging from 380 to 420 bp). The CAG100Het mouse in lane **(B)** exhibited exceptional range of somatic mosaicism for the expanded allele, from 405 bp just above the CAG100 size to 476 bp representing CAG128 size, with clonal drift into a larger allele. The CAG100Het mouse in lane **(C)** displayed less somatic mosaicism with a relatively sharp peak of expanded alleles around 470 bp. The CAG100Hom mouse in the last lane **(D)** showed two expanded alleles of average size 416.65 and 475.64 bp, respectively, again with preferential PCR amplification of the smaller expansion allele.

### Offspring contains less female mutants than expected

In the absence of ATXN2, mouse breeding produces less homozygous mutant and less female pups than expected (4). Also in invertebrates, gender-related reproductive anomalies have been reported, including female sterility in *D. melanogaster* flies with ATXN2 mutations and abnormal masculinization of the germline in *C. elegans* worms with ATX-2 deficiency (36, 37). For these reasons, gender and genotypes were documented among offspring of 25 CAG100Het breeder pairs. The litters contained significantly less homozygous mutants than expected (24% less CAG100Hom than WT pups; p=0.009; *χ*^2^ test with *χ*^2^ = 9.384 and df = 2) and less females than expected (12% reduction; p=0.098; *χ*^2^ test with *χ*^2^ = 2.731 and df = 2) (Table S2). All data suggest that altered ATXN2 functions may impair embryonal development, with some gender-dependence. Thus, the findings constitute evidence for a partial loss-of-function of CAG100-expanded ATXN2 in peripheral tissues and for the high conservation of ATXN2 effects during phylogenesis.

### Initial weight excess reverts over time

We followed the mice over time. The CAG100Hom mice showed progressive motor deficits with hindlimb clasping, reduced strength and ataxia (illustrated in Video S1). Around the age of 14-16 months their motor deficits prompted the veterinarians to sacrifice the animals and prevent suffering. Even before that age, CAG100Hom animals were frequently found dead in their cages without known reasons (*p*<0.001; *χ*^2^=65.366; df= 2; Kaplan-Meier survival analysis with Tarone-Ware test; Fig. 2A).

**Figure 2:**
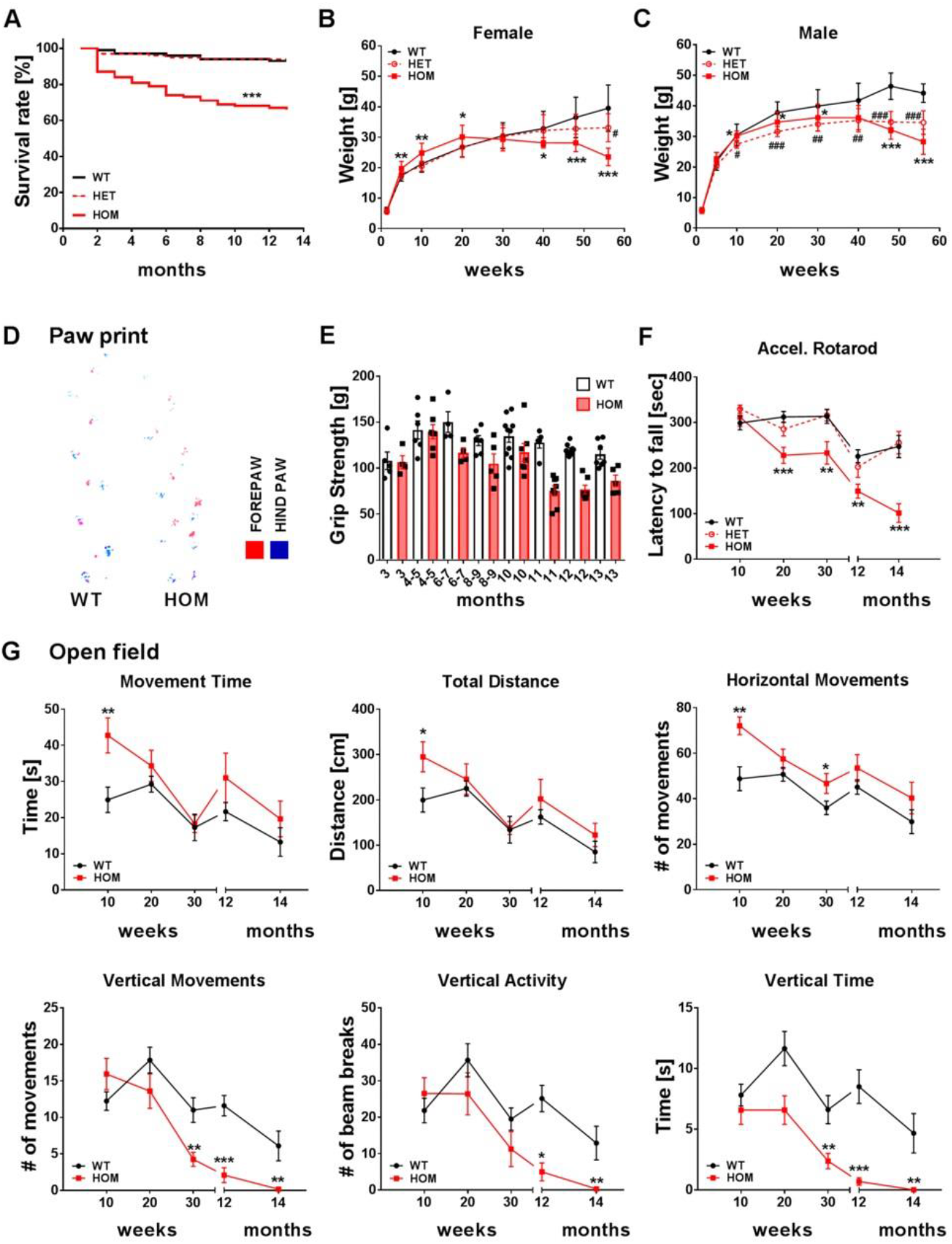
Lifespan, weight and motor phenotypes of *Atxn2*-CAG100-KIN mice across ages. **(A)** The reduction of the ageing cohort through animals found dead at different ages is shown across the lifespan, before all animals were sacrificed for ethical reasons (100% corresponds to 106 CAG100Hom, 235 CAG100Het and 167 WT animals born initially). **(B)** Body weight of female animals and **(C)** of male animals was studied in groups of 4-19 mice per genotype and age. Data is represented as means ± sd. **(D)** Paw prints were recorded in 13 WT and 9 CAG100Hom animals at the age of 12 months. **(E)** Grip strength was assessed in 4-10 animals of each age. **(F)** The latency to fall from a rotarod slowly accelerating from 4 to 40 rpm reflected a very early motor deficit among homozygotes (n = 22 animals per genotype for younger age groups until 8 animals for old ages). **(G)** Open field analyses of the spontaneous movement activity of mice during a 5 minute observation period in an odor-neutral arena recorded in automated manner via infra-red beams for various parameters of locomotion (n = 8-23 animals per age and genotype). Two-way ANOVA; * represents p<0.05, ** p<0.01, *** p<0.001, \*\*\*\**p*<0.0001, \*\*\*\*\**p*<0.00001.

Although all mutant mice eventually developed a loss of weight compared to WT littermates, female CAG100Hom initially gained weight excessively at ages 5-20 weeks. As the disease progressed they lost this extra weight. Female CAG100Het mice also showed reduced body mass during later stages, but without the initial weight gain (p<0.001; F=70.524 with 23 degrees of freedom; two-way ANOVA; Fig. 2B). In male mutants, we did not observe the initial excess in weight, but they also showed a progressive weight loss relative to their WT brothers (*p*<0.001; *F*=121.651 with 23 degrees of freedom; two-way ANOVA; Fig. 2C). In male mutants, weight loss became significant already at the age of 10 weeks in CAG100Het mice and at 20 weeks in CAG100Hom mice. While the homozygous males started to lose weight later than their heterozygous littermates, their weight reduction developed faster and stronger (Fig. 2C).

Thus, all our mutant mice eventually displayed weight loss, but the time course depended on the dosage of expanded ATXN2 and on gender. This is in good agreement with previous reports that homozygous SCA2 patients have a particularly severe disease course (38), so we focused on homozygous animals in further neuropathological and expression analyses. The temporal dynamics of body weight across the lifespan might reflect an initial partial loss-of-function phenotype due to the reduced levels and insolubility of expanded ATXN2, followed by the progressive accumulation in cytosolic aggregates with consequent gain-of-function phenotypes.

### Initial hyperactivity disappears during ageing, later progressive motor deficits are compatible with spinocerebellar ataxia

To establish whether the newly created CAG100Hom mice displayed motor deficits compatible with symptoms observed in human SCA2 patients, we conducted a series of behavioral tests in older mice. Footprint analyses were performed for free movements in a dark tunnel towards the exit, which showed the 12-month-old CAG100Hom mice to display irregular steps (Fig. 2D). Evaluating the particularly vulnerable motor neurons, grip strength analyses were done in CAG100Hom animals from the age of 3 months onward, which revealed a significant decrease in the maximal forelimb efforts over time (*p*<0.001; *F*=10.219 with 7 degrees of freedom; ANOVA). Around the age of 11 months, the forelimb grip strength of CAG100Hom mice became significantly less, while that in WT littermates remained intact (*p*<0.001; *F*=9.964 with 15 degrees of freedom; two-way ANOVA; Fig. 2E). Tests of the motor coordination ability and tenacity to stay on a rotating rod upon slow acceleration showed a significant and stable deficit in CAG100Hom mice from the age of 20 weeks to 12 months, which progressed rapidly at 14 months. CAG100Het animals appeared normal on the rotarod (*p*<0.001; *F*=13.871 with 14 degrees of freedom; two-way ANOVA with Tukey’s post-hoc tests: WT vs. CAG100Hom: p<0.001; WT vs. CAG100Het: p=0.978; Fig. 2F). Assessing spontaneous activity and various motor impairments by open field tests, an initial period of significant hyperactivity around the age of 10 weeks was observed for the parameters movement time, total distance travelled and horizontal movements in CAG100Hom mice (Fig. 2G), as previously described also in *Atxn2*-KO mice (4). Significant reductions in the parameters vertical movement, vertical activity and vertical time indicate problems to balance upright at later ages between 30 weeks and 14 months and may be correlated to progressive spinocerebellar ataxia (Fig. 2G).

### CAG100Het mice show a decreased accuracy in their stepping pattern over time

The CAG100Het mice did not show a deficit on the rotarod (Fig. 2F), despite their loss in body weight (Fig. 2B/C). To quantify disease progression in CAG100Het mice, we used a more refined task and followed their locomotor behavior over time on the ErasmusLadder. The ErasmusLadder is composed of alternating higher and lower rungs. Typically, mice prefer to walk on the upper rungs and avoid touching the lower ones (39). After a week of daily trainings (data not shown) we followed the performance of the mice over time. At 36 weeks of age, at the onset of the weekly training session, both WT and CAG100Het mice showed lower rung touches in approximately 2% of their steps. This percentage remained constant for WT mice until the end of the recording sessions at 52 weeks of age (*p*=0.321; *F*=1.253; repeated measures ANOVA with Greenhouse-Geisser correction; Fig. S3A). In contrast, the heterozygotes gradually became less accurate in their stepping pattern, more than doubling the lower rung touches over time (*p*<0.001; *F*=64.640; two-way ANOVA).

A similar test for the CAG100Hom mice failed, as they proved to be over-sensitive to the pressurized air indicating the trial timing, hampering performance measurements on the ErasmusLadder at the age of 7 months.

### Early symptomatic CAG100Hom show deficient balance and coordination while walking

Since older CAG100Hom mice could not be reliably measured on the ErasmusLadder, we characterized their locomotor phenotype using high-speed videography. To this end, mice were allowed to walk freely on the LocoMouse, essentially consisting of a glass plate between two shelter boxes (40). During the early symptomatic phase, between 7 and 8 months of age, these homozygotes walked slower than the control mice (*p*<0.001; Kolmogorov-Smirnov test; Fig. S3B/C). Under these experimental conditions, mice preferred to walk using a trotting gait, during which the diagonal limbs follow the same pattern (41). The synchrony between starting points of the swing phase of the diagonal limbs was decreased in CAG100Hom mice (*p*<0.001; Kolmogorov-Smirnov test; Fig. S3D). In addition to the slower and less well coordinated trotting pattern, the homozygotes showed signs of improper balance (Video S2/S3). Like patients (42), CAG100Hom had a wide based gait (inter-limb distance: front: WT: 13.9 ± 2.0 mm; CAG100Hom: 17.9 ± 1.7 mm mean ± standard deviation; Z=1.971; *p*=0.049; Z test; Fig. S3B/E). Another sign of impaired balance was the position of the tail, which is largely used for balance control in mice (43). The homozygotes held their tails differently, lifting them further than the WT control mice (*p*<0.001; Kolmogorov-Smirnov test; Fig. S3B/F). Thus, even in the absence of experimental challenges, early symptomatic CAG100Hom at ages of 36-52 weeks showed typical cerebellar signs of slower and less-coordinated walking patterns as well as of balance problems.

### Compensatory eye movements in CAG100Hom mice

The integrity of oculomotor behavior in early symptomatic CAG100Hom mice was tested using two different reflexes promoting gaze stability: the opto-kinetic reflex (OKR), depending on visual input, and the vestibulo-ocular reflex (VOR), depending on vestibular input, combined in the visually-enhanced VOR (VVOR). All reflexes were tested by presenting head-fixed mice with combined sinusoidal visual and/or vestibular input. While the OKR was not affected by the polyQ expansion (*p*=0.281; *F*=1.365; repeated measures ANOVA), the VOR and the VVOR were reduced (*p*=0.025; *F*=8.061 and *p*=0.038; *F*=6.547, respectively; Fig. S3G), in line with the balance problems during locomotion (Fig. S3E/F). Plasticity of the VOR, which critically depends on the cerebellum (44–46), was tested using VOR gain increase training. The mice were trained by visual feedback to make larger eye movements in response to vestibular input, using out of phase vestibular and visual stimulation. Before, in between and after six training sessions of five minutes each, the change in response was evaluated by recording the VOR in the dark. CAG100Hom mice were significantly impaired in this VOR gain-increase adaptation (*p*=0.006; *F*=14.679; Fig. S3H), signified by a clearly reduced change in gain at ages of 7-8 months in these mutants (*p*=0.023; *t*=2.893; *t*-test; Fig. S3H inset).

### Rhythmic behavior impaired in CAG100Hom mice

Licking is a rhythmic behavior in many animals including mice (47–49). Tongue movements are thought to be under control of a central pattern generator, probably located in the hypoglossal zone of the intermediate reticular formation (50), but their rhythm might be modulated by cerebellar activity (51). Like wild-type mice, early symptomatic CAG100Hom animals show rhythmic licking behavior (Fig. S3I). However, the mutant mice displayed a lower licking frequency than the control mice at ages of 7-8 months (*p*<0.001; Kolmogorov-Smirnov test; Fig. S3J-K). Thus, upon careful examination some balance and motor problems can already be detected in early symptomatic CAG100Hom mice.

### Progressive brain atrophy and neuronal aggregation

Analysis of the mutant brain revealed atrophy and weight loss for both sexes in CAG100Hom mice, and to a lesser extent also in CAG100Het mice, at the advanced age of 14 months (Fig. 3A-D). Investigations to detect ATXN2 by immunohistochemistry in the CAG100Hom brain with a monoclonal antibody revealed high signals in many neuron populations, particularly in specific brainstem nuclei (inferior olive and pons), cerebellum, ventral forebrain areas, cerebral cortex and hippocampus, in good agreement with publically available *in-situ* hybridization data of unaffected mice at the Allen Brain Atlas. As a novel observation, some ATXN2 expression was also detectable in oligodendrocytes, e.g. of the corpus callosum (Fig. S4).

**Figure 3:**
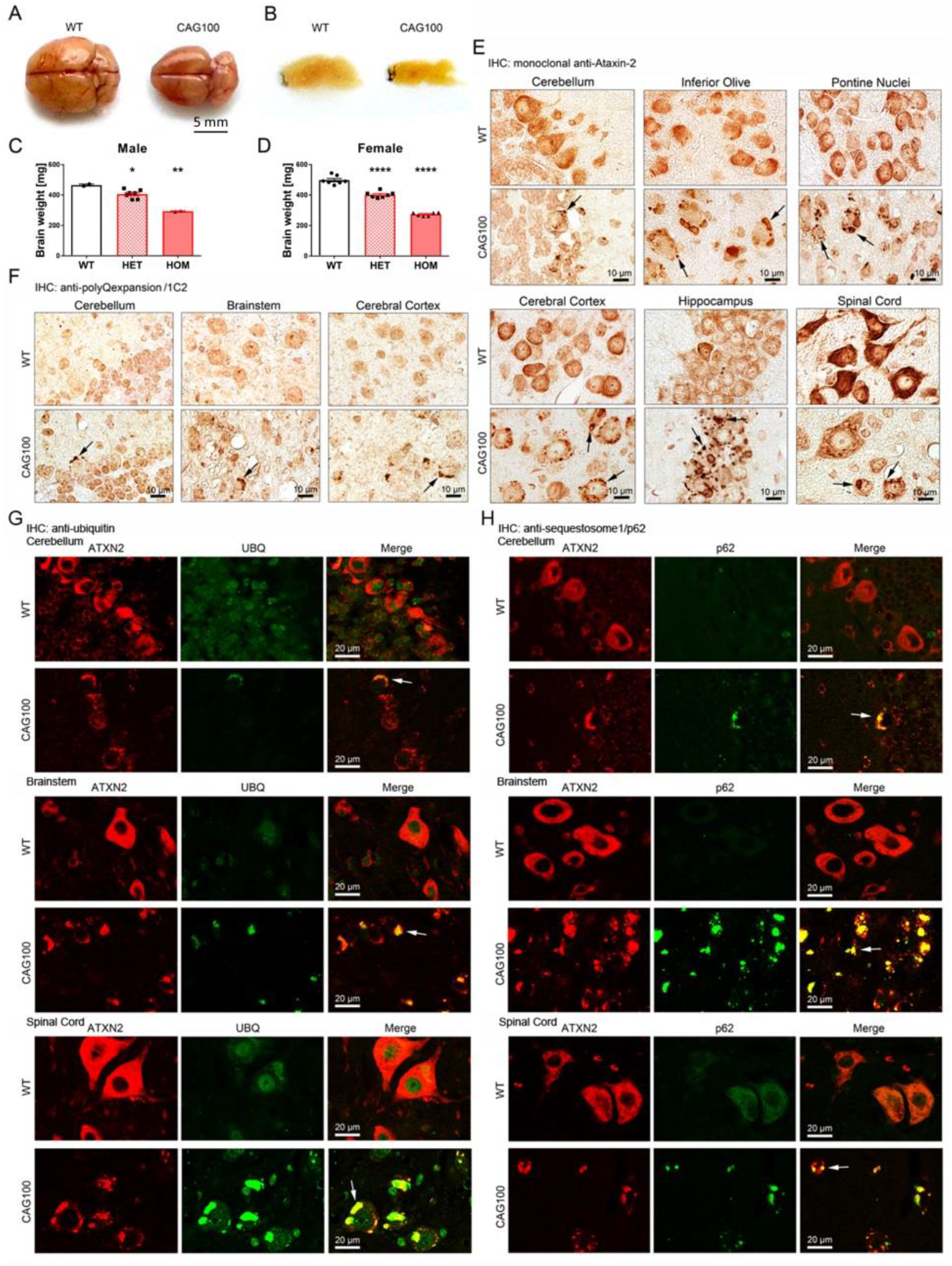
Brain pathology of CAG100Hom mice at the age of 14 months. **(A)** Representative brain photos are shown after animal dissection (seen from above) and **(B)** after dehydration and embedding in paraffin (as sagittal section). Statistical evaluation of brain weight for WT, heterozygous (HET), and homozygous (HOM) animals was compared separately for male **(C)** and female **(D)** animals (for males, reduction to 0.81% vs. 0.55%, p=0.03 vs. 0.001; for females, reduction to 0.87% vs. 0.63%, p<0.00001 vs. <0.00001; tested by ANOVA with multiple testing correction after Bonferroni). Immunohistochemical (IHC) visualization of punctuate or large aggregates, using paraffin-embedded sections and staining for **(E)** ATXN2 with the monoclonal antibody using DAB detection across various brain regions with known SCA2 pathology and strong *Atxn2* expression (black arrows point to cytosolic aggregates), **(F)** polyQ expansion domains with the monoclonal 1C2 antibody using DAB detection, **(G)** co-localization of ubiquitin (UBQ) signals and ATXN2 aggregates using double immunofluorescence (white arrows point to yellow co-localization at aggregates), **(H)** co-localization of the p62 adaptor of ubiquitinated protein aggregates and the ATXN2 signals by double immunofluorescence. Motor neurons and brainstem neurons were more affected by protein aggregates than cerebellar Purkinje neurons.

Electron microscopy of cerebellar Purkinje neurons detected cytosolic protein aggregates (black arrowheads in Fig. S5). In further studies by light microscopy, cytosolic aggregates of ATXN2 were observed in the typical regions affected by neurodegeneration in SCA2, such as cerebellar Purkinje neurons, inferior olivary neurons and pontine nuclei neurons (Fig. 3E upper rows). Aggregates were also detectable in cerebral cortical and hippocampal neurons and in spinal cord motor neurons, where they were particularly large (Fig. 3E lower rows). In all these regions, the cytosolic aggregates were also detectable by immunohistochemistry with the monoclonal anti-polyQ antibody 1C2 (Fig. 3F). Double immunofluorescence was able to show the co-localization of ATXN2-positive cytosolic aggregates with ubiquitin signals (Fig. 3G) and p62 signals (Fig. 3H) in cerebellum, brainstem and spinal cord, suggesting that they undergo the classical elimination via autophago-lysosomal pathways. Again, particularly large protein aggregates could be observed in spinal cord motor neurons (Fig. 3G, lowest row), in good agreement with the preferential vulnerability of motor neurons in pre-symptomatic stages of human SCA2 (52). Thus, this *Atxn2*-CAG100-KIN mouse is an authentic model of the spatio-temporal pattern of pathology known from human SCA2.

### Myelinated axon deficits rather than somatodendritic atrophy explain brain weight loss

To identify the cellular and molecular mechanisms that underlie the selective loss of specific neural circuits, we first focused on the cerebellum in view of the early and progressive ataxia in these animals. Histological sections were stained for fluorescence microscopy, with anti-Calbindin-1 (CALB1) as marker for Purkinje neurons. While brain autopsies from SCA2 patients show prominent neuron loss after 5-25 years with the disease, and other rodent SCA2 models used overexpression to maximize cell death within the 2-year lifespan of mice (28, 32), in our KIN model the loss of neuronal cell bodies and dendritic trees was minor upon histological quantification of Purkinje neuron number and molecular layer thickness. This was true even at final stages when the animals displayed severe motor impairment and had to be sacrificed. This observation was validated in cerebellar homogenates by immunoblots for the somatodendritic marker CALB1 and for the dendritic microtubule component TUBA4A, which is responsible for ALS22. Remarkably, the proteins showed only marginal reductions (CALB1 to 87% ± 0.037 mean ± sem, p=0.031; TUBA4A to 90% ± 0.028, p=0.160), although their transcripts upon qPCR analysis showed almost two-fold deficits (*Calb1* to 57% ± 0.028, p=0.001; *Tuba4a* to 56% ± 0.009, p=0.0001) (Fig. S6). Histological analysis confirmed that Purkinje neurons in CAG100Hom mice at the age of 14 months, while being largely intact, showed reduced expression of CALB1 in a mosaic pattern. In addition, the loss and dispersal of myelinated axons was detectable (Fig. S7C,D). This pathology of myelinated axons was similarly observed also by myelin stains of SCA2 patient cerebellum (Fig. S7A,B). Evident thinning of the deep cerebellar layers, where climbing and mossy fibers arrive from the brainstem and where Purkinje axons relay the efferent signals, was detected upon myelin-specific staining (Fig. S7E,F) and upon hematoxylin-eosin histochemistry (Fig. S7G,H).

Further immunohistochemical analysis with anti-neurofilament antibodies (Fig. S8) observed a slightly diminished axonal neurofilament basket around the Purkinje neuron somata, as well as decreased amounts and an irregular pattern of axons in the lower molecular layer around primary Purkinje dendrites, probably reflecting diminished climbing fiber input from the inferior olivary nucleus of the brainstem at this stage of pathology. In addition, the axons in the granular layer had an irregular distribution, perhaps compatible with abnormal sprouting of mossy fiber input to the cerebellum or of Purkinje axonal output.

### Identification of molecular correlates for axon and myelin loss at protein level

We carried out surveys of cerebellar tissues via global proteomics with label-free mass spectrometry plus MaxQuant / Perseus bioinformatics as well as via global transcriptomics using Affymetrix Clariom D arrays. They confirmed strong and significant depletion of several key factors of axons and myelin in spite of only marginally altered CALB1 levels and relatively preserved dendritic markers in 14-month-old CAG100Hom mice. While these expression profiles suggested multiple pathways of pathogenesis and have to be evaluated fully with more samples at diverse ages in future manuscripts, we used significant findings from these screenings as a basis for further molecular studies. In the present manuscript they were focused on the validation of consistent downregulations, concerning key protein components of neuronal axons and of oligodendrocyte myelin, employing additional samples with independent methods.

First, essential factors of axonal projections were quantified by immunoblots in cerebellar homogenates, demonstrating their deficiency by normalizations relative to beta-Actin (ACTB) as general tissue housekeeping protein (Fig. 4). To assess their preferential vulnerability in comparison to the Purkinje neuron somatodendritic compartment, additional normalizations versus CALB1 were also shown. Reductions were marked for the axonal cytoskeleton factors NEFH (Neurofilament Heavy Chain, diminished to 63% ± 0.107, upon normalization to CALB1, p=0.046 upon t-test without multiple testing correction), NEFL (Neurofilament Light Chain, 68% ± 0.090, p=0.085) and of the synaptic cell adhesion factor NPTN (Neuroplastin, 78% ± 0.038, p=0.006), but not for NEFM (Neurofilament Medium Chain, 99% ± 0.115, p=0.997). Similarly marked reductions at the transcript level were detected for *Nefh/Nefm/Nefl* (40% ± 0.015, p=0.0002; 46% ± 0.016, p=0.0005; 48% ± 0.024, p=0.0003, respectively) and *Nptn* (71% ± 0.015, p=0.001) (see Fig. 4).

**Figure 4:**
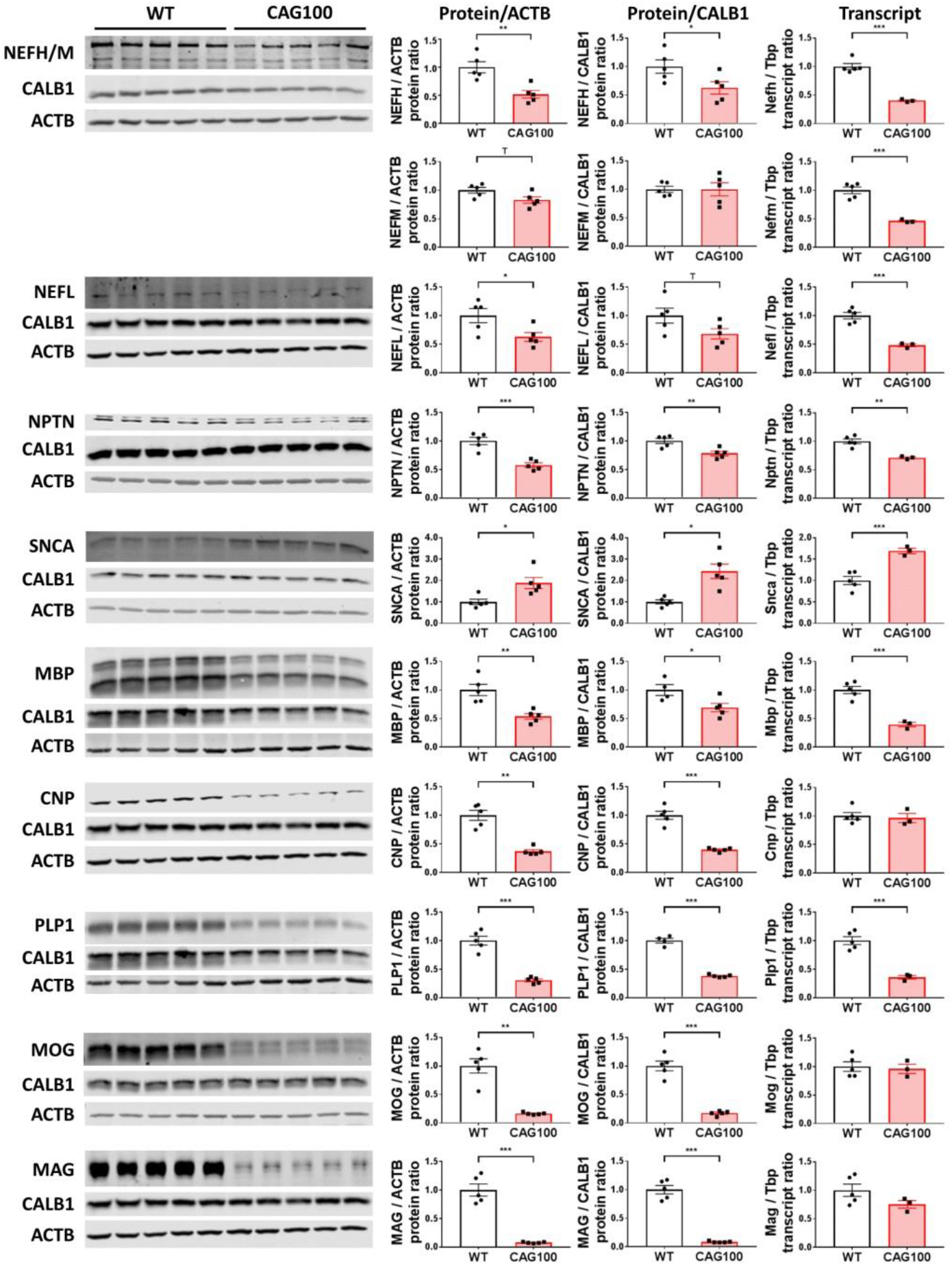
Protein and mRNA dysregulation in cerebella of CAG100Hom mice at the age of 14 months. Dysregulations of protein and mRNA levels were observed for several factors involved in axon myelination, namely the axon-localized Neurofilament heavy / medium / light chain subunits (NEFH / NEFM / NEFL), the synapse-localized Neuroplastin (NPTN), the synapse-localized alpha-Synuclein (SNCA), versus the oligodendrocyte-localized Myelin Basic Protein (MBP), 2’,3’-Cyclic Nucleotide 3’ Phosphodiesterase (CNP), Proteolipid Protein (PLP1), Myelin oligodendrocyte glycoprotein (MOG) and Myelin associated glycoprotein (MAG). The abundance of the protein of interest was normalized versus beta-Actin (ACTB) as total tissue loading control, and separately versus Calbindin-1 (CALB1) as marker of Purkinje neuron somatodendritic compartment preservation. The mRNA abundance of each factor is shown with normalization versus TATA-box-binding protein (*Tbp*) mRNA, to elucidate if the dysregulation might be triggered at the transcriptional level by the RNA-binding ATXN2 expansion via direct interaction, or represents a post-transcriptional event. T-test was used in general.

The downregulation did not affect all axonal and presynaptic components uniformly, given that alpha-synuclein showed strong upregulations at protein (to 243 ± 0.341, p=0.012) and mRNA level (to 169% ± 0.062, p=0.0008) (Fig. 4). Alpha-synuclein is a component of the SNARE complex that associates with neurotransmitter vesicles in the presynaptic compartment (53) and inhibits excitatory neurotransmission (54).

Second, substantial downregulations were also observed for the main myelin components MBP (Myelin Basic Protein, reduced to 69% ± 0.072 upon normalization to CALB1, p=0.041), CNP (2’,3’-Cyclic Nucleotide 3’ Phosphodiesterase, 40% ± 0.015, p=0.001), and PLP1 (Proteolipid Protein 1, 38% ± 0.011, p=0.0004). Dramatic depletions were found for the myelin glycoproteins MOG (Myelin Oligodendrocyte Glycoprotein, 17% ± 0.018, p=0.0004) and MAG (Myelin Associated Glycoprotein, 8% ± 0.006, p=0.0002). Again strong decreases at the mRNA level were detected for *Mbp* (to 39% ± 0.038, p=0.0002) and *Plp1* (36% ± 0.033, p=0.0003), but not for *Cnp, Mog* and *Mag* (Fig. 4).

### RNA-binding ATXN2-Q100 affects further myelin and axon factors at mRNA level

In addition, we observed significant reductions in mRNA levels in the 14-month-old CAG100Hom cerebellum of several other factors for which no specific antibody was available with sufficient sensitivity to quantify the endogenous protein. For some transcripts, dysregulations were also found in the Atxn2-KO mice, which were studied at the age of 6 months when their progressive obesity phenotype is already evident. Among the relevant myelin components shown in Fig. 4, a dysregulation in the cerebellum of *Atxn2*-KO mice was not observed for *Cnp* or *Plp1*, but for *Mbp* (KO to 73% ± 0.035 compared to its WT control, p=0.007). Dysregulation in CAG100 versus KO cerebellum was also documented for *Mal* (Myelin And Lymphocyte Protein, CAG100 60% ± 0.031 compared to WT, p=0.001, KO 113% ± 0.069, p=0.129) and *Mobp* (Myelin-Associated Oligodendrocyte Basic Protein) long (CAG100 27% ± 0.027, p=0.0002; KO 71% ± 0.065, p=0.005) versus short transcript (CAG100 39% ± 0.009, p=0.0007, KO 108% ± 0.136, p=0.634) (Fig. 5A).

**Figure 5:**
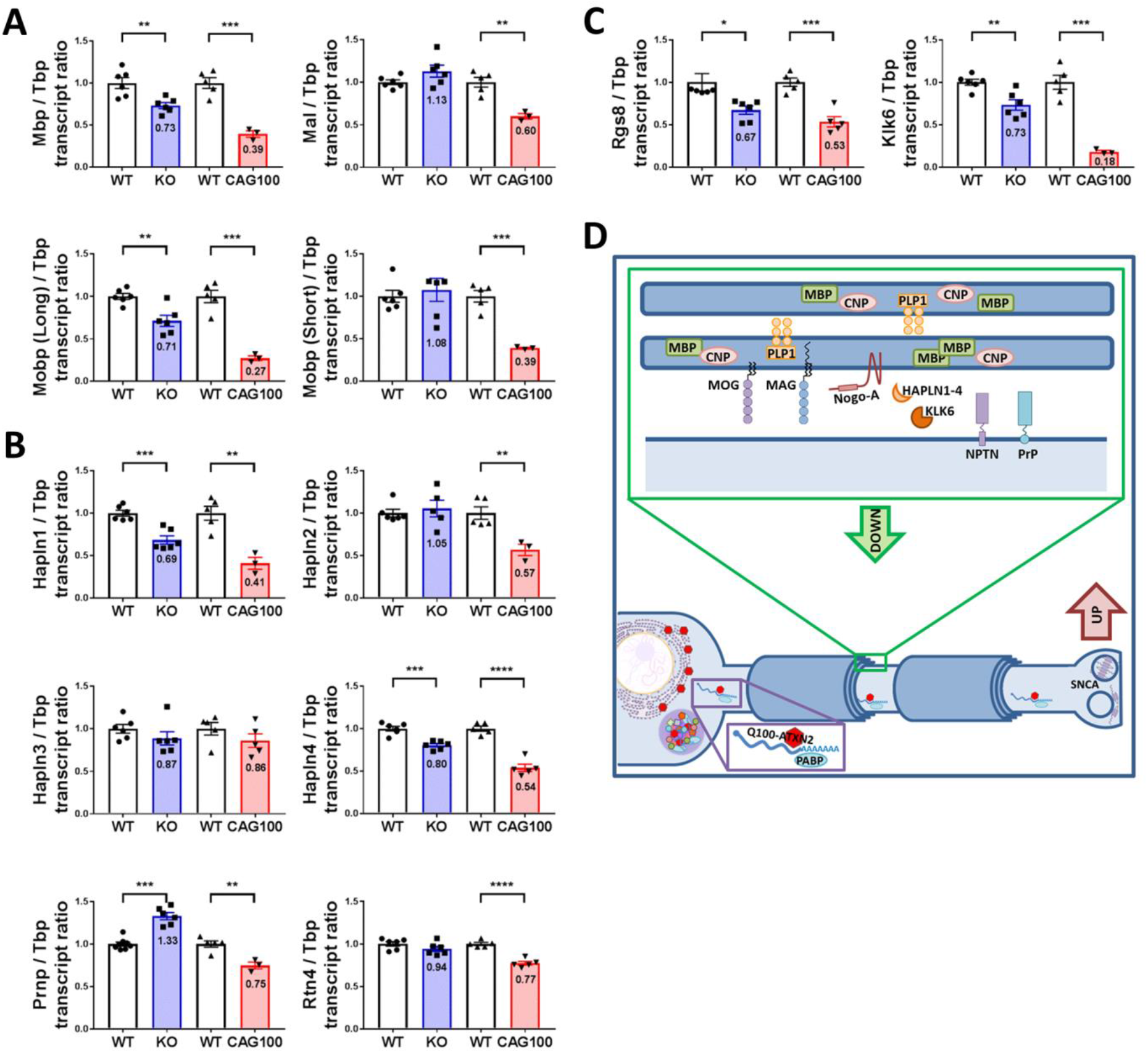
Transcript dysregulations in cerebella of CAG100Hom versus *Atxn2*-KO mice. Downregulations of mRNA levels were observed for several factors involved in axon myelination, where specific and sensitive antibodies were unavailable. **(A)** Among oligodendrocytic factors, this included *Mal* (Myelin and Lymphocyte Protein) and two isoforms of *Mobp* (Myelin-Associated Oligodendrocyte Basic Protein), in addition to *Mbp* (Myelin Basic Protein), which is shown for comparison. **(B)** Among neuronal factors, this included *Hapln1, Hapln2, Hapln3, Hapln4* (Hyaluronan and Proteoglycan Link proteins), *Prnp* (Prion protein) and *Rtn4* (NoGo-A). Only the Prion protein mRNA was dysregulated in the opposite direction in *Atxn2*-KO tissue. **(C)** Among factors of unclear origin and without membrane association, this included *Rgs8* (Regulator of G Protein Signaling 8) and *Klk6* (Kallikrein Related Peptidase 6). Analysis was performed for KO mice at the age of 6 months when obesity is evident, comparing with CAG100Hom mice at the age of 14 months when weight loss is strong. Fold-changes are shown within the bars, t-test was used in general. **(D)** Schematic representation of the dysregulated transcripts and proteins in the cerebellum of 14-month-old CAG100Hom mouse.

Among relevant neuronal components with membrane association, a dysregulation in CAG100 as well as KO cerebellum was observed for components of the extracellular matrix at perineuronal nets and Ranvier nodes, namely *Hapln1* (Hyaluronan and Proteoglycan Link Protein 1, CAG100 to 41% ± 0.070, p=0.002; KO 69% ± 0.049, p=0.0005) and *Hapln4* (CAG100 54% ± 0.043, p<0.0001; KO 80% ± 0.021, p=0.0003) rather than *Hapln2* (CAG100 57% ± 0.067, p=0.005, KO 105% ± 0.098, p=0.640) and *Hapln3* (CAG100 86% ± 0.081, p=0.237, KO 87% ± 0.078, p=0.258), as well as the axonal myelination triggers *Prnp* (Prion Protein, CAG100 75% ± 0.067, p=0.006; KO 133% ± 0.042, p=0.0001) rather than *Rtn4* (Nogo-A, CAG100 77% ± 0.022, p<0.0001, KO 94% ± 0.030, p=0.179) (Fig. 5B).

Among excitability factors that are cytosolic or extracellular, this was also true for *Rgs8* (Regulator of G Protein Signaling 8, CAG100 53% ± 0.060, p=0.0004; KO 67% ± 0.049, p=0.023) as a previously reported dysregulation in SCA2 mouse models, and for *Klk6* (Kallikrein Related Peptidase 6, CAG100 18% ± 0.017, p=0.0004; KO 73% ± 0.061, p=0.005) as a secreted protease implicated in cell adhesion, in degradation of presynaptic alpha-synuclein and in turnover of oligodendroglial MBP (55) (Fig. 5C). Given that ATXN2 is a RNA-binding protein and that the above mRNAs involved in axon myelination are prominently affected, this pathway may be crucial at the early stage of cerebellar atrophy in SCA2. Significant dysregulation of the factors schematically depicted in Fig. 5D, and the axonal disconnection probably explains the movement deficits at ages when no neuronal loss is detectable.

## Discussion

We generated a genetic mouse model of SCA2, which faithfully reflects the spatial distribution of affected neural pathways, with the preferential vulnerability of motor neurons, vestibular and cerebellar pathways resulting in chronically progressive locomotor deficits. These phenotypes appear to be triggered by axonal disconnection, demyelination and neural dysfunction rather than by loss of neural cell bodies within the lifespan of our mouse mutants. As knock-in model it reflects the partial loss-of-function of ATXN2 in peripheral proliferating tissues and at the initial stage of neural pathology, while neurons at older age are stressed by the progressive toxicity of ATXN2 due to the formation of protein aggregates.

This partial loss-of-function is well explained by its strong insolubility in neural tissue and its reduced abundance in peripheral cells. Similar findings of reduced levels and insolubility of polyQ expanded disease proteins were also reported for SCA7 (56). For polyQ expanded Ataxin-3 it was shown that expression and solubility was normal in induced pluripotent stem cells, fibroblasts or glia cells, but changed to an insoluble aggregated state upon neuronal differentiation and exposure to excitatory stimuli such as glutamate (34). These previous observations explain our findings that ATXN2-Q100 is quite soluble in fibroblasts, although in much lower amounts, while it goes into immediate insolubility and aggregation in neural tissue. As consequences of this partial loss of function, we observed reduced numbers of female mutants among the offspring, weight excess and hyperactivity during early life. Interestingly, the changes in body weight show an initial increase followed by progressive decrease not only in our mouse model, but also in SCA2 families upon careful longitudinal assessment (57).

The progressive loss of body weight and brain weight is compatible with an insidious increase of ATXN2-Q100 toxicity due to aggregate formation. ATXN2 is expressed in pancreas and affects the islet beta-cells in their trophic state and their insulin secretion (4), so we assume that ATXN2-Q100 aggregate toxicity affects these postmitotic cells via the known effects of ATXN2 on mTORC1 signaling (7, 58), thus triggering a depletion of body fat stores. Similarly, the strong weight reduction of the brain might be explained in large part by a loss in myelin fat. ATXN2 expansion sizes of Q>100 clearly trigger myelination defects also in man, since these SCA2 patients develop widespread leukoencephalopathy upon brain imaging (59). The importance of demyelination for SCA2 patients with shorter expansions is unclear.

Overall the strongest effect of the CAG100 expansion concerned two glycoproteins that are crucial for myelin-axon adhesion, oligodendroglial MOG and MAG, diminishing their abundance below 20% at the protein level. Importantly, the transcript levels of both factors were not altered, so this alteration could be a downstream consequence of axon-myelin pathology with excessive turnover of MAG / MOG proteins, or it might be a direct consequence of oligodendrocytic ATXN2 dysfunction impairing the translation of these mRNAs at the rough ER. Both MAG and Nogo-A (encoded by *Rtn4*) interact as alternative ligands for the receptor NgR1, which modulates neurite outgrowth (60). Nogo-A was transcriptionally downregulated by the toxic ATXN2-Q100. Conversely, a previous report showed the knock-out mouse of Nogo-A to trigger a transcriptional upregulation of ATXN2 and of Ataxin-1 (61), so this adhesion pathology might be among the important primary events of pathogenesis.

Given that the expression of ATXN2 in oligodendrocytes is low in comparison to neurons, it is important to ask which factors that are intrinsic to the axons themselves are dysregulated by the CAG100 expansion. From the study of peripheral nerves of SCA2 patients it is known that axonal degeneration in sensory afferents is among the first objective signs of disease (62), in agreement with the observations on preferential axonal deficits in our new mouse model. Interestingly, among all factors studied in the cerebellum only the prion protein (*Prnp*) mRNA levels were upregulated with significance by the *Atxn2*-KO as well as conversely downregulated with significance by the CAG100 expansion, suggesting a toxic gain-of-function mechanism. We have previously shown that the deletion of the ATXN2 ortholog PBP1 leads to a downregulation of the prion RNQ1 at the protein level in yeast (12). Prion protein is an axonal membrane glycolipoprotein and controls myelin maintenance signals via the G-protein coupled receptor Adgrg6 (63). It is well known that toxic protein aggregates of the prion protein lead to a spongiform encephalopathy in man, cows, sheep and mice (64).

The CAG100 expansion also had a marked effect on the mRNA and protein levels of the synaptic adhesion factor neuroplastin (NPTN), which is important for the localization of GABA_a_ and GluA1 neurotransmitter receptors (65). A converse 2-fold upregulation at the protein level and 1.5-fold upregulation at the transcript level was noted for the synaptic vesicle release inhibitor alpha-synuclein. Gene triplication or duplication events leading to increased 1.5 to 2-fold dosage of alpha-synuclein were shown to cause Parkinson’s disease at manifestation ages around 35 and 55 years, respectively (66), via the accumulation, fibrillation and aggregation of insoluble alpha-synuclein in so-called Lewy bodies (67, 68). In view of ATXN2 expansions also causing levodopa-responsive Parkinson’s disease by unknown molecular mechanisms, this induction of alpha-synuclein may contribute to pathogenesis in SCA2 or modify it. In addition, the CAG100 expansion led to downregulations of *Hapln1, Hapln2* and *Hapln4*, three extracellular factors that are concentrated in perineuronal nets and at axonal Ranvier nodes between myelinated segments, where they control the diffusion of cations that are crucial for saltatory conductivity (69, 70). As components of the axonal cytoskeleton, the neurofilament chain subunits NEFH/NEFM/NEFL showed marked reductions at the protein and mRNA level, indicating either strong loss or shrunk diameter of axons.

As mentioned above, only *Prnp* expression reflected the loss versus toxic gain of function in ATXN2, while *Mbp, Mobp, Hapln1, Hapln4, Rgs8* and *Klk6* mRNA dysregulations showed downregulations both in *Atxn2*-KO and in *Atxn2*-CAG100-KIN cerebella, with the stronger effect always occurring in the KIN tissue (Figure 7). Reflecting an excessive rather than partial loss-of-function of ATXN2 in CAG100 tissue, these data are compatible with the notion that compensatory changes of the Ataxin-2-like protein (ATXN2L) and other interactor molecules such as PABPC1 within the same pathway are able to limit downstream effects in the KO tissue, whereas in the KIN tissue the aggregation of ATXN2 triggers a sequestration of its interactor molecules into insolubility and a non-compensated loss-of-function for the entire pathway.

The preferential affection of cell-cell adhesion factors and membrane proteins by the CAG100 expansion might be explained by several aspects of the physiological function of ATXN2. Firstly, ATXN2 is normally localized at the rough ER rather than smooth ER or free polysomes (6), so it seems to modulate the mRNA translation of factors that are trafficked to the membranes or the extracellular space. Secondly, ATXN2 modulates the availability of glucose, fatty acids and amino acids (10), so it may modify also the synthesis of glycolipoproteins and of myelin fat. Thirdly, at stress granules ATXN2 is important for the quality control of RNAs that were damaged during cytosolic transport, an event that will occur with high probability for factors that undergo extensive trafficking from the nucleus along the whole axon length to synaptic adhesion sites.

Overall, the resulting neural disconnection and degeneration affects the nervous system with cytosolic inclusion bodies of ATXN2 in the characteristic pattern of olivo-ponto-cerebellar atrophy (OPCA), as was carefully documented in patients (71, 72). The aggregates are particularly large in spinal motor neurons in our mouse model, and indeed it was recently shown in SCA2 mutation carriers that motor neuron degeneration appears even before the onset of cerebellar ataxia, accompanied by muscle cramps, impaired conduction velocity due to axon demyelination and the loss of subcutaneous fat tissue (52, 73, 74).

As a result of all these molecular changes, motor deficits develop over time. In previous SCA2 mouse models, performance on the rotarod typically deteriorated over time. The longer the polyQ repeat, the earlier such effects could be observed: as early as 8 weeks for transgenic mice with a long Q128-repeat (29) to as late as 18 months for knock-in mice with a Q42-repeat (33) with onset in-between for repeat sizes midway (14, 28). Shorter repeats (Q22) did not develop a phenotype on the rotarod (32). Smaller steps and decreased performance on the beam walk have also been reported (14, 28). To the best of our knowledge, other types of behavior have not been tested before on mouse models for SCA2. Our behavioral data confirms the aforementioned earlier reports on other SCA2 mouse models regarding the deficits on the rotarod, the stepping pattern and the beam walk. The locomotor phenotype is in line with cerebellar deficits (39, 40). The grip strength test points towards motor problems, while the intact OKR shows that visual input is not (strongly) affected during the early stage of the disease. Remarkably, the tasks involving the vestibular system showed early deficits, pointing towards dysfunction of the vestibulo-cerebellar system as one of the first hallmarks in the disease process.

In conclusion, the traditional view of SCA2 as a disorder of neural cell loss in the typical OPCA distribution has to be complemented with novel evidence on the prominent vulnerability of spinal motor neurons and the precocious atrophy of axons with their myelin, via dysregulation of adhesion factors. The subsequent neural disconnection underlies the movement deficit, which progresses insidiously from the age of 6 months to a maximal lifespan of 16 months, and triggers the characteristic features of vestibulo-cerebellar and oculomotor dysfunction.

## Methods

### Body Weight and Behavioral Observations

Heterozygous *Atxn2*-CAG100-KIN mice were used for breeding. Offspring with CAG100Hom and WT or CAG100Het genotype of similar ages and identical sex were used as case-control pairs for phenotypic comparisons. Sudden death of animals was noted together with the age at death. Mice were weighed before behavioral testing. In contrast to all other measurements, male and female animals were separated for weight analyses due to strong gender-specific weight differences. Brain weight was measured after cervical dislocation, dissection and removal of the olfactory bulb, employing an analytical balance. If not otherwise stated, male as well as female animals were used for phenotype studies without separation. Grip strength was assessed by measuring the peak force of the fore limbs in 10 trials per mouse on an electronic grip strength meter (TSE, Bad Homburg). Paw prints were evaluated by painting the forepaws with a non-toxic red ink, the hind limbs of mice with blue. The mice were placed at one end of a dark tunnel, so that their walk to the other end will leave paw prints on the white paper that covers the floor (tunnel 6 cm high × 9 cm wide × 40 cm long). Footprint movement patterns were analyzed as described previously (33). Assessment on an accelerating rotarod apparatus (model 7650 Robert & Jones, Ugo Basile, Comerio) and in an open field arena (Versamax, Omnitech, Columbus, Ohio) occurred as previously described (33). During the acceleration of the rotarod from 4-40 rpm, every mouse had four consecutive 6 minute trials interrupted by at least 10 min of break, without previous training. The latency to fall was recorded for each trial; the mean value of the four trails was calculated and used for statistical analysis. Videorecording occurred at ages from 10 to 14 months. For the beam test, the animals had to walk across a surface with length of ∼1 m and a diameter of 18 mm. For the clasping test mice were suspended by their tails for about 1 min. Behavioral analyses were always conducted at the same daytime to avoid variances caused by circadian clock.

### Perfusion

Mice were anesthetized with an overdose of Ketaset (300 mg/kg) and Domitor (3 mg/kg) by an intraperitoneal injection. To assess the anesthetic depth the withdrawal reflex was monitored. Intracardial perfusion was done with phosphate buffer saline (PBS) followed by 4% paraformaldehyde (PFA) in 0.1 M PBS. For paraffin embedded sections, the tissue was post-fixed overnight in 4% PFA at 4 °C, dehydrated and incubated in paraplast for 24 h at 56 °C. All tissues were cut and mounted in 7-μm-thick slices using a microtome. For cryosections, the tissue was also post-fixed overnight in 4% PFA at 4 °C, immersed in 30% sucrose until it sank, cut with a cryostat in 30-μm-thick slices and kept in cryoprotection solution (30% ethylene glycol, 25% glycerin, 0,01% sodium azide in 0.1 M PBS) at −20 °C until used.

### RNA Isolation and Expression Analysis

Whole brain was removed after cervical dislocation; cerebellum and two hemispheres were dissected into separate tubes and immediately frozen in liquid nitrogen. RNA extraction from cerebellum was performed with TRIzol Reagent (Invitrogen) according to user manual. DNase digestion was performed using DNaseI Amplification Grade (Invitrogen), and cDNA synthesis from 1 μg of total RNA template was performed by the SuperScript IV VILO kit (ThermoFisher) according to manufacturer’s instructions. To assess the gene expression changes, quantitative real-time PCR analyses were performed with StepOnePlus Real-Time PCR System (Applied Biosystems) equipment. cDNA from 25ng total RNA was used for each PCR reaction with 1 μl TaqMan^®^ Assay, 10 μl FastStart Universal Probe Master 2x (Rox) Mix and ddH_2_O up to 20 μl of total volume. The TaqMan^®^ Assays utilized for this study are: *Atxn2* (Mm01199894_m1), *Calb1* (Mm00486647-m1), *Cnp* (Mm01306641_m1), *Hapln1* (Mm00618325_m1), *Hapln2* (Mm00480745_m1), *Hapln3* (Mm00724203_m1), *Hapln4* (Mm00625974_m1), *Hprt1* (Mm00446968_m1), *Klk6* (Mm00478322_m1), *Mag* (Mm00487538_m1), *Mal* (Mm01339780_m1), *Mbp* (Mm01266402_m1), *Mobp* long transcripts (Mm02745649_m1), *Mobp* short transcripts (Mm01348317_g1), *Mog* (Mm00447824_m 1), *Nefh* (Mm01191456_m1), *Nefl* (Mm01315666_m1), *Nefm* (Mm00456201_m1), *Nptn* (Mm00485990_m1), *Plp1* (Mm01297210_m1), *Prnp* (Mm004483 89_m 1), *Rgs8* (Mm01290239_m1), *Rtn4* (Mm00445861_m1), *Snca* (Mm00447333_m1), *Tbp* (Mm00446973_m1), *Tuba4a* (Mm00849767_s1). The PCR conditions were 50 °C for 2min, 95 °C for 10 min, followed by 40 cycles of 95 °C for 15 s and 60 °C for 1 min. Gene expression data was analyzed using 2^-ΔΔCt^ method (75) with *Tbp* or *Hprt1* as housekeeping genes.

### Protein extraction and Western blots

Cerebellar tissue was homogenized with a motor pestle in 5-10x weight/volume amount of either RIPA buffer [50 mM Tris-HCl (pH 8.0), 150 mM NaCl, 2 mM EDTA, 1% Igepal CA-630 (Sigma), 0.5% sodium deoxycholate, 0.1% SDS and one tablet Complete Protease Inhibitor Cocktail (Roche)] or LD buffer [low detergent buffer: 20 mM Tris-HCl (pH 8.0), 137 mM NaCl, 1% glycerol, 0.1% NP-40, 2 mM EDTA]. Following centrifugation, the pellets were dissolved in SDS lysis buffer [137 mM Tris-HCl (pH 6.8), 4% SDS, 20% glycerol, and one tablet Complete Protease Inhibitor Cocktail (Roche)] in order to obtain insoluble proteins. Cell pellets from MEF cultures were homogenized in 150 μl RIPA buffer. Protein concentration was determined with a Spectrophotometer (Eppendorf) using 5x Bradford Reagent (Roti-Quant, Carl Roth). 20 μg of total proteins were mixed with 2x loading buffer [250 mM Tris-HCl pH 7.4, 20% Glycerol, 4% SDS, 10% 2-Mercaptoethanol, 0.005% Bromophenol blue, 5% dH_2_O], incubated at 90 °C for 2 min, separated on polyacrylamide gels and were transferred to Nitrocellulose membranes (GE Healthcare). The membranes were blocked in 5% BSA/TBS-T, and incubated overnight at 4 °C with primary antibodies against 1C2 (Chemicon #MAB1574), ATXN2 (monoclonal from BD Biosciences #611378, 1:500; polyclonal from Proteintech #21776-1-AP, 1:500), ACTB (Sigma #A5441, 1:10000), CALB1 (Cell Signaling #13176, 1:2500), CNP (Cell Signaling #5664S, 1:1000), MAG (Cell Signaling #9043S, 1:500), MBP (Merck #05-675, 1:250), MOG (Abcam #ab32760, 1:500), NEFH/M (Proteintech #18934-1-AP, 1:500), NEFL (Cell Signaling #2837S, 1:1000), NPTN (Alomone Labs #ANR-090, 1:500), PLP1 (Abcam #ab28486, 1:1000), SNCA (Cell Signaling #2642S, 1:1000), TUBA4A (Aviva Systems #ARP40179-P050, 1:500). Fluorescent labeled secondary goat anti-mouse (IRDye 800CW, Li-Cor, 1:10000) and goat anti-rabbit (IRDye 680RD, Li-Cor, 1:10000) antibodies were incubated for 1 h at room temperature (RT). Membranes were visualized using Li-Cor Odyssey Classic instrument. The image analysis to quantify signal intensities was performed using ImageStudio software.

### Immunocytochemistry, immunohistochemistry and histological stains

For immunocytochemistry, 1×10^5^ cells from WT and CAG100Hom MEF cultures were seeded on PDL coated 12 mm cover slips. 24 h later, the cells were washed and stressed with 0.5 mM NaARS supplemented in the DMEM growth medium for 45 min at 37 °C. Control cells were washed and supplemented with only DMEM growth medium for 45 min. Cells were washed once before fixation with 4% paraformaldehyde/PBS at RT for 20 min, then were permeabilized with 0.1% Triton-X-100/PBS for 20 min at RT. Blocking was done with 3% BSA/PBS solution for 1 h at RT. Primary antibody incubation with PABPC1 (Abcam ab21060, 1:100) and ATXN2 (BD Biosciences #611378, 1:100) antibodies was performed in 3% BSA/PBS for 1 h at RT. Secondary antibody incubation with goat anti-rabbit-Alexa Fluor 546 (Molecular Probes, 1:1000), goat anti-mouse-Alexa Fluor 488 (Molecular Probes, 1:1000) antibodies and DAPI was performed in 3% BSA/PBS for 1 h at RT in dark. The coverslips were mounted on glass slides with fluorescent mounting medium and dried overnight. Cell imaging was performed using Zeiss LSM 510 using a Plan Apochromat 100 × objective microscope, and ImageJ software was used to merge images.

For immunohistochemistry, paraffin embedded sections were rehydrated in a descending alcohol series. Bull’s Eye Decloaker (1:20) was used for antigen retrieval and the sections were incubated with the following primary antibodies overnight: anti-1C2 (Millipore #MAB1574, 1:800), anti-ATXN2 (BD Bioscience #611378, 1:50), anti-p62 (Santa Cruz #sc25575, 1:50) and anti-Ubiquitin (UBQ, Dako #ZO458, 1:100). For DAB stainings, Vector NovaRED Peroxidase kit was used after blocking the endogenous peroxidase with 100% methanol, 30% H_2_O_2_ in Tris/HCl pH7.6 (1:1:18) for 30 min. For fluorescent stainings, Cy-3 anti-mouse IgG and Cy-2 anti-rabbit IgG (Dainova, 1:1000) were used. Calbindin staining and neurofilament/p62 coimmunofluorescence staining were performed on free floating sagittal cerebellar cryosections. The sections were permeabilized with 0.3% Triton X-100 in 0.1 M PBS for 30 min, blocked with 10% BSA, 0.3% Triton X-100 in 0.1 M PBS for 90 min and incubated with the primary antibodies (24 h at RT for anti-Calbindin-1 from Sigma #C9848, 1:300; overnight at 4 °C for anti-Neurofilament 200 kDa from Chemicon #MAB5256, 1:200 and for anti-p62 from Santa Cruz #sc25575, 1:50) followed by Alexa Fluor 488 and Alexa Fluor 565 secondary antibodies for 2 h at RT. Nuclei were stained with DAPI. Double fluorescence immunohistochemical stainings were studied with the Nikon confocal microscope Eclipse 90i and a 60x magnification. The Leica 090-135-001 microscope was applied for single immunohistochemical stainings with a 60x magnification. For histological myelin stains, human sections were treated with the modified Heidenhain protocol as described before (76), for the mouse sections the Hito Luxol Fast Blue – PAS OptimStain PreKit (Hitobiotec Corp, Kingsport, TN) was used. Hematoxylin & Eosin staining was done according to standard protocols. The SCA2 patient (biobank# GR 245-07) analyzed by myelin stain was male with CAG repeat sizes 39/20 in the *ATXN2* gene, disease onset at 32 years of age, death at 51 years, the control individual was age- and sex-matched.

### Statistics

Unless specified otherwise, all statistical tests were performed as unpaired Student’s t-test using GraphPad Prism software version 4.03 (2005) after establishing that each population was normally distributed (one-sided Kolmogorov-Smirnov test), with figures displaying mean values and standard error of the mean (sem), unless stated otherwise. Values p<0.05 were considered significant.

### Study approval

The animal studies were performed with ethical approval of the government office in Darmstadt (FK/1083).

## Author contributions

MVH, NES, JCP, BWK, MSc, ED, KS, UR, DM, MM, PH, CIDZ, SG, LWJB and GA designed the experiments, MVH, NES, JCP, BWK, MSe, ED, KS, DM, MM, LEM and SG performed the experiments, MSe contributed novel analysis tools, MVH, NES, JCP, BWK, MSe, MSc, ED, KS, UR, DM, MM, CIDZ, LEM, SG, LWJB and GA analyzed the data, MVH, NES, MSc, LWJB and GA wrote the text with contributions of all authors, DM, MSc, CIDZ and GA contributed funding.

## Acknowledgements

The authors wish to thank Birgitt Meseck-Selchow and the staff at the Zentrale Forschungs-Einrichtung at Frankfurt University Medical School for their assistance with animal assessments, Nadia Khosravinia, Ilja IJpelaar and Laura Post of the Neuroscience Department at the Erasmus Medical Center for technical support, Dr. Dennis Eckmeier of the Champalimaud Foundation for help with setting up the LocoMouse Tracker software and Beata Lukaszewska-McGreal for technical assistance with the assessment of protein levels.

Financial support was provided by the Deutsche Forschungs-Gemeinschaft (AU 96/ 11-1 and 11-3), the European Research Council Starting Grant (ERC-Stg, 680235) (MSc), the Netherlands Organization for Scientific Research (NWO-ALW; CIDZ), the Dutch Organization for Medical Sciences (ZonMW; CIDZ), Life Sciences (CIDZ), and ERC-adv and ERC-POC (CIDZ), and the Max Planck Society. Michel Mittelbronn would like to thank the Luxembourg National Research Fund (FNR) for the support (FNR PEARL P16/BM/11192868 grant).

## Notes

The authors have declared that no conflict of interest exists.

